# Structure-guided reinterpretation of a disease-associated CWH43 residue-533 truncation identifies internal disruption of a conserved C-terminal module in idiopathic normal pressure hydrocephalus

**DOI:** 10.64898/2026.05.01.722067

**Authors:** Haowei Cao, Zhihan Yan, Jing Wang, Yue Liu, Zheng Wei, Yu Cheng, Shuqun Hu, Dejun Yang

## Abstract

Idiopathic normal pressure hydrocephalus (iNPH) is a chronic neurological disorder characterized by ventricular enlargement together with gait impairment, cognitive decline, and urinary dysfunction. Disease-associated CWH43 variants have emerged as genetic clues in iNPH, but the molecular consequences of truncating variants remain poorly defined. Here, we used structure-guided analysis to reinterpret the disease-associated human CWH43 variant c.1596del, annotated as p.Leu533Ter and referred to here as the residue-533 truncation. AlphaFold-based modeling of the full-length 699-amino-acid protein supported a conserved C-terminal D3 module spanning residues 407-699. Within this framework, the residue-533 truncation is best interpreted as an internal interruption of D3 rather than deletion of a poorly constrained terminal tail. Pocket analysis identified a dominant candidate surface site within D3 containing 18 pocket-lining residues, 55.6% of which (10/18) are removed by truncation. Independent cavity mapping further showed that the affected region is better understood as a broader cavity-bearing architecture rather than a single local pocket. Cross-boundary contact analysis, multi-model reproducibility, and representative local geometry supported structural coupling between retained and deleted D3 segments, indicating that truncation is predicted to cause retained-side decoupling in addition to direct residue loss. Together, these findings support a model in which the disease-associated residue-533 truncation disrupts a structurally integrated C-terminal module and its associated cavity-bearing architecture. This framework refines interpretation of CWH43-associated iNPH variants and provides a mechanistic bridge between human genetic association and architecture-level loss of function.

## Introduction

Idiopathic normal pressure hydrocephalus (iNPH) is a chronic disorder of aging classically characterized by ventriculomegaly together with progressive gait impairment, cognitive decline, and urinary dysfunction. Although the syndrome can be partially shunt-responsive, diagnosis remains challenging because clinical and radiologic findings overlap with other age-related neurologic disorders, and not all patients improve after cerebrospinal fluid diversion. Increasing evidence further suggests that iNPH is not merely a nonspecific disorder of CSF circulation, but a biologically heterogeneous disease in which inherited risk, epithelial and ciliary dysfunction, and membrane-trafficking pathways may all contribute to pathogenesis (Carswell 2023; Johnson and Williams 2025; Mehta et al. 2025; Nakajima et al. 2021; Pearce et al. 2024; Räsänen et al. 2024).

Among the emerging genetic signals, CWH43 has drawn particular interest. Initial work implicated CWH43 deletions in iNPH, and subsequent cohort-based studies identified additional truncating and missense CWH43 variants, supporting CWH43 as a mechanistically informative candidate gene. At the same time, later population-based analyses indicated lower variant prevalence in some cohorts, suggesting that the contribution of CWH43 may be cohort-or population-dependent rather than a uniformly distributed high-penetrance signal (Piccinin et al. 2025; Räsänen et al. 2024; Räsänen et al. 2025; Tipton et al. 2023; Yang et al. 2021).

The biological function of mammalian CWH43, however, remains incompletely defined. In yeast, Cwh43 is required for remodeling the lipid moiety of glycosylphosphatidylinositol (GPI) anchors toward ceramide-containing species, whereas in mammalian systems the broader PGAP/GPI remodeling pathway is known to regulate endoplasmic reticulum exit, Golgi maturation, stable cell-surface expression, and membrane-raft association of GPI-anchored proteins. Prior overexpression studies using GFP-tagged mammalian CWH43 in HeLa cells further suggested association with the ER/Golgi membrane system, consistent with a role in membrane maturation and GPI-anchor-related trafficking, although these observations are best interpreted as supportive localization evidence rather than definitive endogenous localization (Yang et al. 2021). Related work on PGAP2, PGAP5, downstream GPI-anchor remodeling, and recent GPI biosynthesis reviews therefore provides an important conceptual framework for interpreting how CWH43 dysfunction could disturb membrane organization, trafficking, and cell-surface protein homeostasis even if the exact biochemical activity of mammalian CWH43 is not yet fully established (Fujita et al. 2009; Fujita et al. 2011; Guo et al. 2024; Kinoshita and Fujita 2016; Maeda et al. 2007; Tashima et al. 2006; Umemura et al. 2007).

This framework is supported by prior experimental work on CWH43-associated hydrocephalus models. In mice, Cwh43 loss was reported to cause hydrocephalus, gait and balance abnormalities, reduced ependymal cilia, and altered localization of GPI-anchored proteins at the apical surfaces of choroid plexus and ependymal cells, linking CWH43 deficiency to ventricular pathology and epithelial surface organization in vivo. These observations strengthen the rationale for treating disease-associated CWH43 variants not only as genetic markers of risk but also as mechanistic entry points into membrane maturation and CSF-barrier biology (Yang et al. 2021).

Within this context, the human CWH43 variant c.1596del (ENST00000226432.9, MANE Select), annotated as p.Leu533Ter, is of particular interest. Hereafter, we refer to this lesion mechanistically as the residue-533 truncation. By creating a premature termination codon (PTC), this lesion may have both transcript-level and protein-level consequences. It has been reported in multiple iNPH-associated cohorts and provides a tractable opportunity for structure-guided mechanistic analysis. Rather than representing a simple distal shortening event, this lesion raises a more specific structural question: whether truncation at residue 533 disrupts a conserved C-terminal module from within. Addressing this question is especially important because such an internal-truncation model would provide a more informative bridge between human genetic association and architecture-level loss of function than a generic “missing tail” interpretation.

## Results

### Residue-533 truncation is an internal truncation within the conserved C-terminal D3 module

The main mechanistic dissection presented below was performed on the full-length rat Cwh43 structural framework, which provided the most complete context for module-level mapping of the residue-533 truncation (Fig. 1a; Supplementary Fig. S1a). Human-side analyses were used as an architecture-level confirmation layer and to anchor the manuscript-level interpretive framework around the disease-associated variant p.Leu533Ter (Fig. 1d; Supplementary Fig. S1d-f; Supplementary Table S2). Although the human-side work was not expanded into a second full pocket/cavity workflow, inspection of the human full-length model together with InterPro-based annotation supported the same high-level conclusion: residue 533 lies within a domain-supported structured C-terminal region rather than a distal unstructured tail (Fig. 1d; Supplementary Fig. S1d-f).

**Figure 1.**
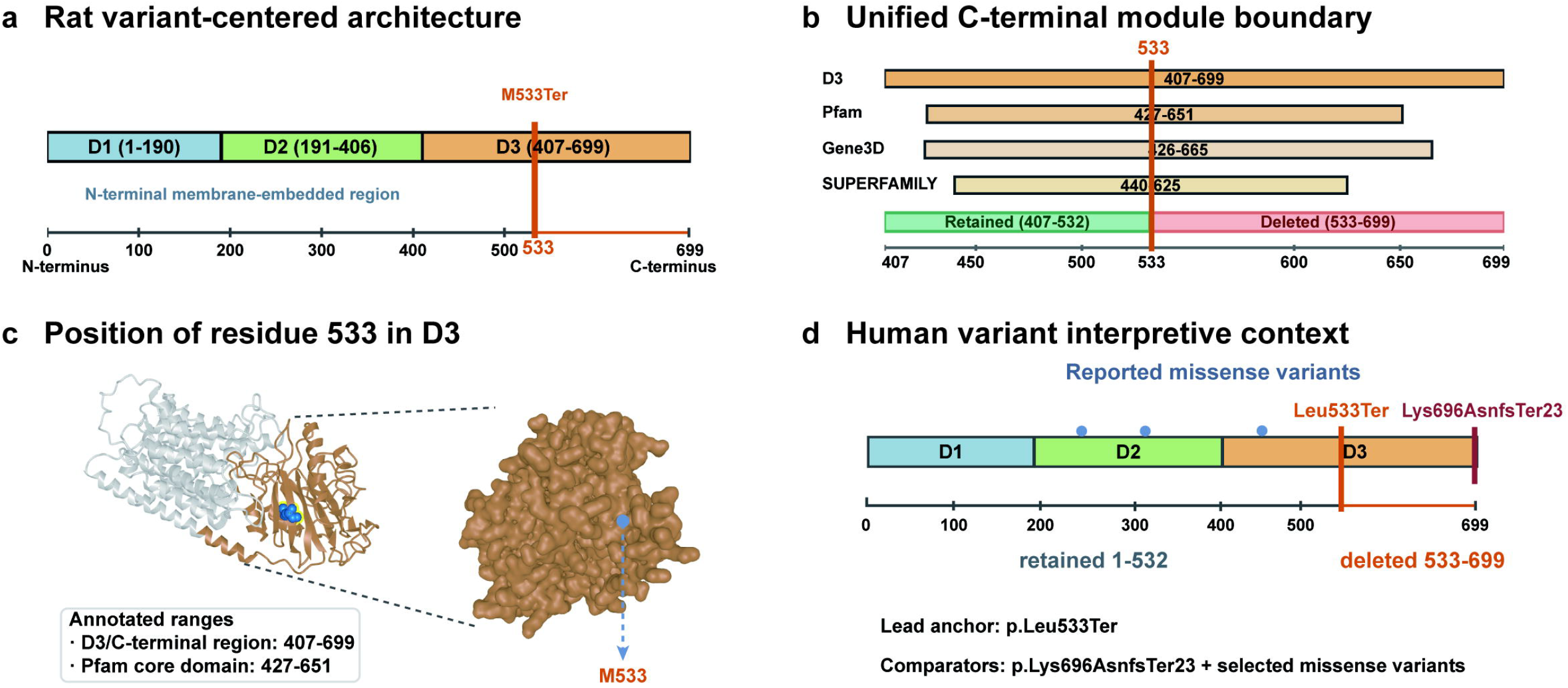
Disease-associated residue-533 truncation is internal to the conserved C-terminal module a, Rat variant-centered full-length structural framework used for mechanistic dissection, showing the D1–D3 architecture and the position of residue 533 within the C-terminal D3 module. b, Rat C-terminal boundary annotation showing overlap between the D3 interval and the Pfam-, Gene3D-, and SUPERFAMILY-supported conserved C-terminal region; the residue-533 truncation boundary falls within this structured module rather than beyond it. c, Minimal three-dimensional placement of M533 within the rat D3 scaffold, illustrating that the site lies within a folded C-terminal region rather than at a distal terminal edge. d, Human variant interpretive context, highlighting human CWH43 c.1596del (ENST00000226432.9, MANE Select), p.Leu533Ter as the lead human truncation anchor and retaining p.Lys696AsnfsTer23 together with selected reported missense variants as human interpretive-context comparators. The residue-533 position falls within a domain-supported structured C-terminal region in the human full-length context. Human-side domain support is provided by InterPro-based annotation of the C-terminal region, whereas fuller architectural confirmation and module-level transferability support are provided in Supplementary Fig. S1d–f.

AlphaFold modeling of the full-length 699-aa protein yielded a highly convergent, high-confidence structural framework, with a pTM of 0.94, a ranking score of 0.95, and a low disordered fraction of 0.02, supporting use of the predicted fold for domain-level interpretation (Fig. 1a; Supplementary Fig. S1a). Within this framework, the C-terminal D3 region forms a structured module rather than a distal terminal remnant, and the residue-533 boundary falls within the same C-terminal region highlighted by Pfam-, Gene3D-, and SUPERFAMILY-based support (Fig. 1b; Supplementary Fig. S1b; Supplementary Fig. S2a).

Cross-species multiple-sequence alignment of the D3 region, performed using MAFFT and Clustal Omega during iterative optimization and cross-checking, converged on the same qualitative conclusion: D3 is the most evolutionarily constrained C-terminal module in the current working architecture (Fig. 1b; Supplementary Fig. S1f; Supplementary Fig. S2b). In parallel, rat-anchored regional support and hotspot-framing analyses placed the residue-533 boundary within the same structured C-terminal D3/Pfam-supported core selected for downstream structural dissection (Supplementary Fig. S2a-d).

Together, these observations support interpreting the residue-533 truncation as an internal truncation within a domain-supported, evolutionarily constrained C-terminal module rather than as a minor terminal shortening event (Fig. 1a-c; Supplementary Fig. S1a-f). Inspection of the full-length human model, together with human truncation framing and cross-species transferability support, extends this interpretation to the disease-associated human residue-533 context at the module level rather than by assuming residue-wise identity (Fig. 1d; Supplementary Fig. S1d-f; Supplementary Table S2). Even without assigning a specific biochemical activity to D3, the available evidence indicates that residue 533 is mechanistically consequential because the truncation interrupts a coherent C-terminal structural unit from within (Fig. 1; Supplementary Fig. S1). This conservation framing is mechanistically important because the residue-533 lesion does not truncate a loosely constrained terminal segment, but interrupts an evolutionarily constrained C-terminal scaffold from within. In this study, conservation is therefore most informative at the module and regional level, where it helps distinguish internal module disruption from a generic tail-loss model.

### Cross-boundary structural coupling explains retained-side decoupling after residue-533 truncation

Consistent with the internal-truncation interpretation established in Fig. 1, the wild-type D3 structure contained a dense cross-boundary contact network linking residues retained N-terminal to the residue-533 boundary with residues deleted C-terminal to the boundary (Fig. 2a). These contacts indicate that the deleted C-terminal segment is not loosely appended but structurally integrated with the retained D3 fragment through boundary-spanning packing relationships; the residue-level components of this retained/deleted architecture are summarized in Supplementary Table S1.

**Figure 2.**
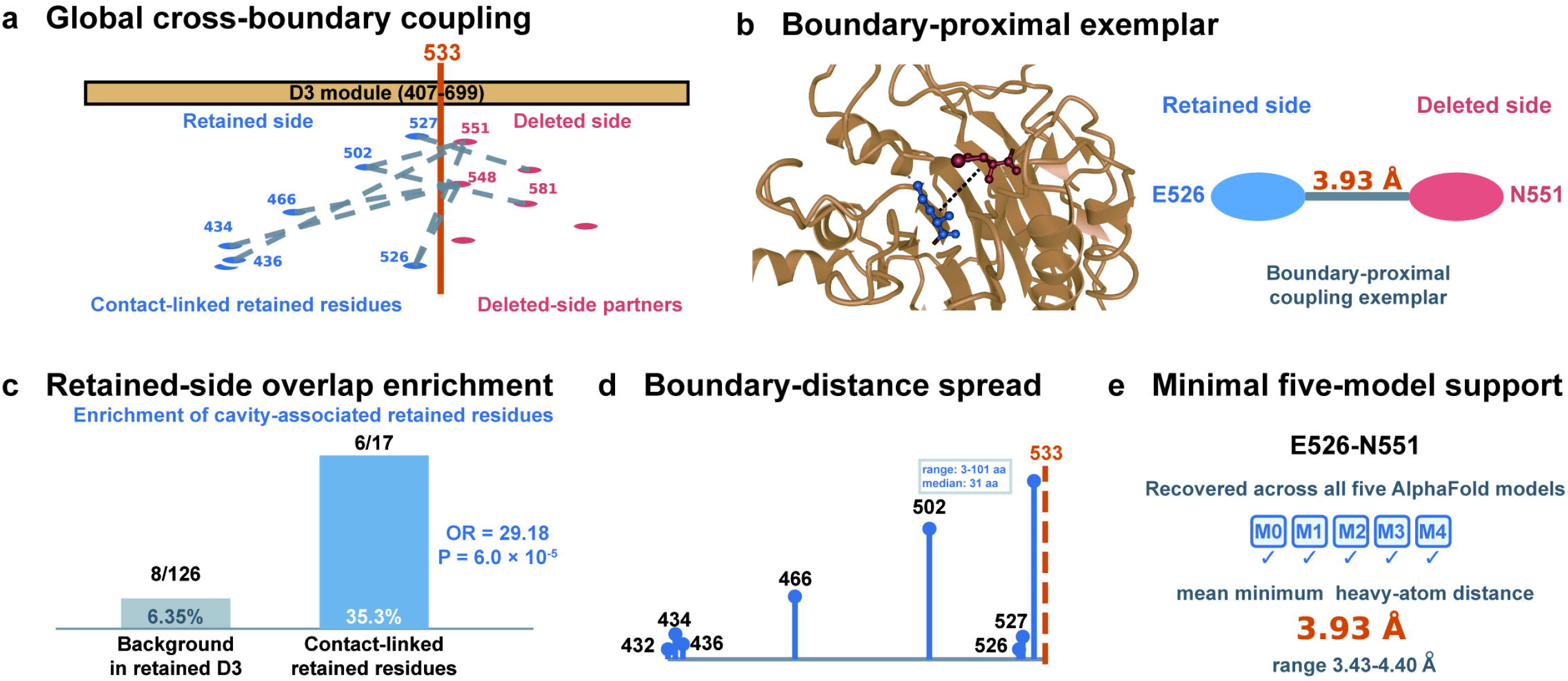
Residue-533 truncation disrupts cross-boundary structural coupling within D3 **a**, Global D3 coupling map showing that residues retained N-terminal to the residue-533 boundary are structurally linked to residues deleted C-terminal to the boundary, indicating that the deleted segment is integrated with the retained D3 fragment rather than loosely appended. **b**, Boundary-proximal coupling exemplar showing a local structural view of retained-side E526 and deleted-side N551 within the wild-type D3 scaffold, together with a simplified right-side summary of the corresponding mean minimum heavy-atom distance across five AlphaFold models. **c**, Retained-side overlap enrichment analysis. At a 6.0 Å heavy-atom contact threshold, 17 retained-side residues contacted deleted-side Pocket 1 residues, and 6/17 of these contact-linked retained residues were themselves retained Pocket 1 residues, compared with a background frequency of 8/126 across the retained D3 segment (odds ratio = 29.18, P = 6.0 × 10^-5). **d**, Boundary-distance spread of contact-linked retained residues. Retained-side residues contacting deleted-side pocket-associated partners spanned positions 432–530, corresponding to 3–101 residues from the residue-533 boundary (median = 31 aa), indicating that the retained-side effect extends beyond the immediate truncation edge. **e**, Five-model reproducibility support showing that the boundary-proximal E526–N551 coupling relationship is recovered across all five AlphaFold models, arguing against interpretation as a single-model artifact. Together, these results support a model in which residue-533 truncation destabilizes the retained D3 fragment not only through direct residue deletion but also through loss of cross-boundary stabilizing contacts required for native local packing.

Representative local geometry supported this interpretation. In the wild-type D3 scaffold, retained-side E526 was spatially proximate to deleted-side N551, providing a boundary-proximal exemplar of cross-boundary coupling (Fig. 2b; Supplementary Table S1). Across all five AlphaFold models, the mean minimum heavy-atom distance for this pair was reproducibly short (3.93 ± 0.35 Å, range 3.43–4.40 Å), supporting persistent local packing across the residue-533 boundary (Fig. 2e; Supplementary Fig. S3a, f).

Quantitative overlap analysis further indicated that the retained-side structural effect was not randomly distributed across the preserved D3 fragment. At a 6.0 Å heavy-atom contact threshold, 17 retained-side residues contacted deleted-side Pocket 1 residues, and 6/17 of these contact-linked retained residues were themselves retained Pocket 1 residues, compared with a background frequency of 8/126 across the retained D3 segment (odds ratio = 29.18, P = 6.0 × 10^-5; Fig. 2c). This enrichment localizes retained-side decoupling preferentially to cavity-associated retained residues rather than to the retained segment as a whole, consistent with the residue-class framework and quantitative support layers shown in Supplementary Fig. S3b, c and catalogued in Supplementary Table S1.

Boundary-distance analysis showed that these retained-side contact-linked residues were not confined to the immediate truncation edge but instead spanned positions 432–530, corresponding to distances of 3–101 residues from the residue-533 boundary (median = 31 aa; Fig. 2d). Five-model comparison further showed that the boundary-proximal E526–N551 coupling relationship was recovered across all five AlphaFold models (Fig. 2e; Supplementary Fig. S3a, f), indicating that the observed cross-boundary organization is reproducible and unlikely to reflect a single-model peculiarity. Together, these data support a model in which residue-533 truncation destabilizes the retained D3 fragment not only through direct residue deletion but also through loss of cross-boundary stabilizing contacts required for native local packing (Fig. 2a–e; Supplementary Fig. S3a–d; Supplementary Table S1).

### Residue-533 truncation disrupts the dominant predicted surface pocket within D3

Pocket analysis of the isolated D3 module (residues 407–699), performed using a PrankWeb/P2Rank-based workflow, identified the dominant predicted pocket recovered in this region, designated Pocket 1 (Fig. 3a). This site received a P2Rank score of 13.44 with an associated probability of 0.646 and comprised 18 pocket-lining residues, making it the strongest pocket candidate in the isolated D3 region despite only modest pocket-level conservation (Fig. 3a; Supplementary Fig. S2b–d).

**Figure 3.**
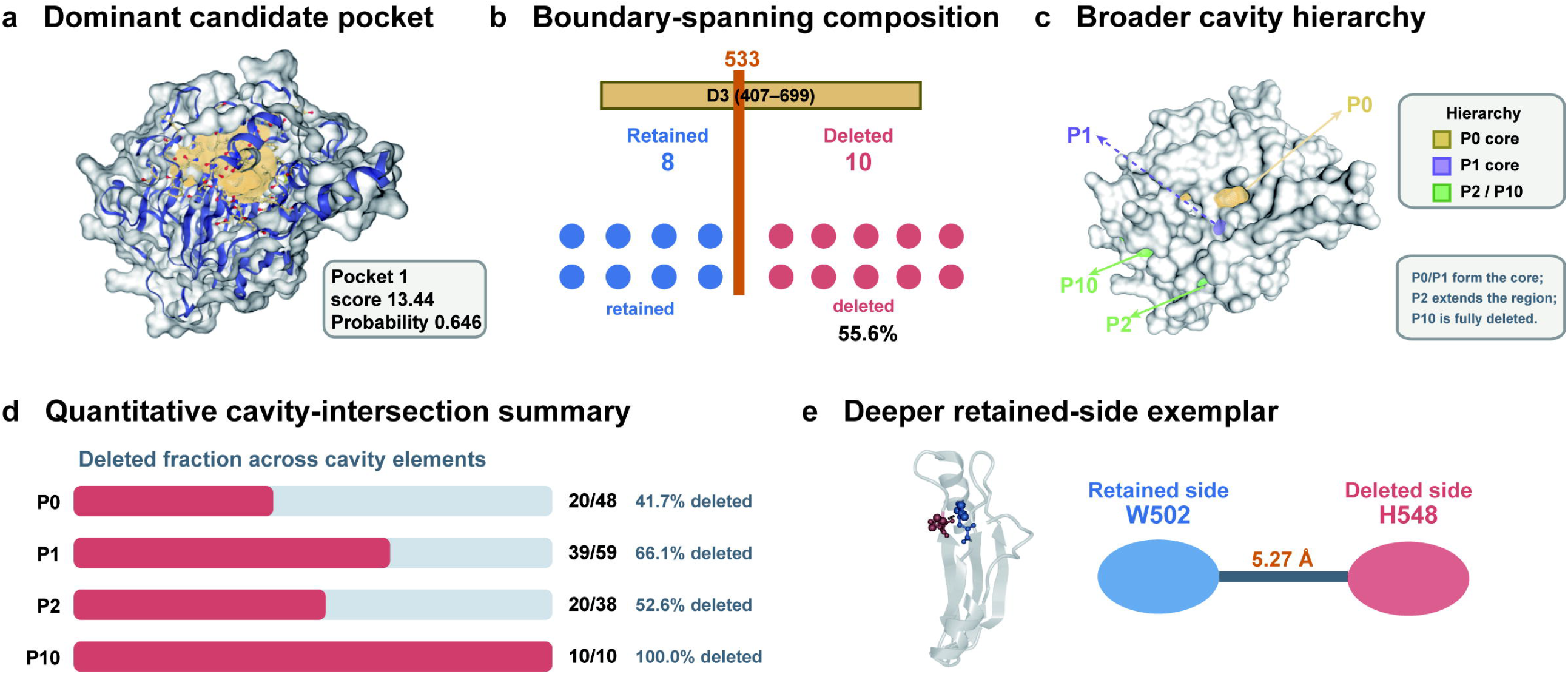
Residue-533 truncation disrupts a candidate pocket within a broader cavity-bearing C-terminal module **a**, Dominant candidate pocket. PrankWeb/P2Rank analysis of the isolated D3 region identified one dominant predicted pocket (Pocket 1), with a score of 13.44 and an associated probability of 0.646. **b**, Boundary-spanning composition of Pocket 1. Pocket 1 contains 18 pocket-lining residues and spans the residue-533 truncation boundary, with 8 residues retained and 10 residues deleted after truncation. **c**, Broader cavity hierarchy. Independent cavity mapping resolves a broader cavity-bearing architecture in D3, comprising two core components (P0 and P1), an extension pocket (P2), and a smaller secondary component (P10). **d**, Quantitative cavity-intersection summary. Residue-level mapping showed that 20/48 P0 residues, 39/59 P1 residues, 20/38 P2 residues, and 10/10 P10 residues fall C-terminal to residue 533 and would therefore be deleted by truncation. **e**, Deeper retained-side exemplar showing a local structural view of retained-side W502 and deleted-side H548 within the cavity-associated region of the wild-type D3 scaffold, together with a simplified right-side summary of the corresponding mean minimum heavy-atom distance across five AlphaFold models. Together, these panels indicate that residue-533 truncation is predicted to disrupt not only a dominant candidate pocket, but a broader cavity-bearing structural unit.

Pocket 1 spans residues 434, 436, 437, 466, 474, 502, 526, 527, 548, 550, 551, 552, 581, 583, 611, 612, 635, and 636 (Fig. 3b; Supplementary Table S1). Only 8 of these 18 residues lie within the retained segment upstream of the truncation boundary, whereas 10 of 18 residues, corresponding to 55.6% of the pocket, fall C-terminal to residue 533 and are therefore deleted by the residue-533 truncation (Fig. 3b). This distribution indicates that the predicted pocket crosses the truncation boundary rather than residing entirely within the preserved N-terminal fragment of D3 (Fig. 3b; Supplementary Table S1).

These data support a structural interpretation in which the residue-533 truncation is expected to substantially disrupt, rather than merely weaken, the dominant candidate surface site in D3 (Fig. 3a, b). Although the pocket-level conservation signal was modest, the site remained structurally credible because it was recovered from the isolated D3 region, embedded within a broader high-confidence local framework, and situated within the same rat-anchored D3/Pfam-supported core highlighted in Supplementary Fig. S2a-d. Importantly, the modest pocket-level conservation signal does not negate the broader structural inference, because Pocket 1 is embedded within the more strongly supported D3/Pfam-defined conserved region intersected by the truncation boundary.

### Independent cavity mapping supports a broader cavity-bearing architecture in D3

Independent cavity analysis using a DoGSiteScorer-style workflow indicated that the structurally relevant feature in D3 is better understood as a broader cavity-bearing region than as a single isolated pocket (Fig. 3c). Residue mapping identified two major adjacent pockets (P0 and P1), a neighboring extension pocket (P2), and a smaller secondary pocket (P10), together forming a connected cavity-associated architecture intersected by the truncation boundary (Fig. 3c; Supplementary Table S1).

Residue-level analysis further showed that these DoGSite-defined pockets are substantially intersected by the residue-533 truncation. P0 comprised 48 lining residues, of which 20/48 (41.7%) mapped to residues C-terminal to 533 and would therefore be deleted by truncation; P1 comprised 59 lining residues, of which 39/59 (66.1%) fell in the deleted segment; P2 comprised 38 lining residues, with 20/38 (52.6%) deleted by truncation; and P10 comprised 10 lining residues, all of which (10/10; 100%) were located C-terminal to residue 533 (Fig. 3d). Thus, the truncation is predicted to affect not only one local cavity element but multiple pockets contributing to a larger C-terminal surface region (Fig. 3c, d; Supplementary Table S1).

Comparison with the previously defined PrankWeb Pocket 1 supported the same interpretation because the PrankWeb pocket did not map exclusively to a single DoGSite pocket but instead overlapped with both P0 and P1, more strongly with P1 than with P0 (Fig. 3a–d). Representative deeper retained-side geometry further supported this broader structural-consequence model: in the wild-type D3 scaffold, retained-side W502 was spatially proximate to deleted-side H548 within the cavity-associated region (Fig. 3e; Supplementary Fig. S3f). Across five AlphaFold models, the mean minimum heavy-atom distance for this pair was 5.27 Å, indicating that cavity-associated packing relationships are predicted to be lost not only immediately at the truncation boundary but also at deeper retained-side positions within D3. The same logic is reinforced by deeper supplementary exemplars, lightweight network support, and the integrated geometry table (Supplementary Fig. S3d–f; Supplementary Table S1). Taken together, these results support a model in which the D3 region contains a broader cavity-bearing architecture that is structurally intersected by the residue-533 truncation boundary and is therefore disrupted at the module level rather than only at the level of a single local pocket (Fig. 3a–e; Supplementary Fig. S3c–f; Supplementary Table S1).

## Discussion

The main value of the present study is not that it defines a definitive biochemical mechanism for CWH43, but that it substantially narrows the mechanistic landscape for a disease-associated residue-533 truncating variant. The most defensible conclusion from the current structure-guided analyses is that the residue-533 truncation should not be interpreted as a minor distal shortening event. Rather, it is best understood as an internal interruption of a conserved C-terminal structural unit, with consequences extending from direct residue loss to broader disruption of local architectural organization. In this respect, the present work refines interpretation of a human disease-associated CWH43 truncation from a generic loss-of-function label to a more specific module-level structural disruption model.

Several lines of evidence converge on this interpretation. Residue 533 maps within a structured and evolutionarily constrained C-terminal region rather than within a loosely organized terminal tail. The dominant predicted D3 surface pocket is intersected directly by the truncation boundary, with 10 of 18 pocket-lining residues removed, while independent cavity mapping indicates that the relevant structural feature in this region is better understood as a broader cavity-bearing architecture rather than as a single narrow pocket. In parallel, cross-boundary contact analysis, enrichment of cavity-associated retained residues among deleted-side contact partners, boundary-distance spread, and representative local geometry support the view that truncation is predicted to induce retained-side decoupling in addition to direct residue loss. Collectively, these results argue against a simple ‘missing tail’ interpretation and instead support disruption of a structurally integrated C-terminal module whose native organization depends on interactions spanning the truncation boundary. In this study, conservation is most informative at the module and regional level: the key point is that residue 533 falls within an evolutionarily constrained C-terminal framework, so truncation disrupts part of a conserved structural scaffold rather than merely deleting a poorly constrained tail segment.

Population-level interpretation, however, still requires caution. Although the present structure-guided analyses support a severe architecture-level consequence for the residue-533 truncation within the conserved C-terminal module, this does not by itself imply a fully penetrant Mendelian effect in all carriers. A more plausible interpretation is that this lesion acts within a context-dependent risk framework, in which genetic background, aging-related factors, and tissue-specific vulnerability modulate phenotypic expression. In this view, structural severity and population-level tolerance are not necessarily contradictory, but may instead reflect reduced penetrance, late manifestation, or dependence on additional biological cofactors. This perspective is also consistent with the broader clinical heterogeneity of iNPH and reinforces the importance of distinguishing architecture-level loss-of-function inference from deterministic disease prediction.

At the same time, the present study has clear limits. The dominant D3 surface feature identified here should be regarded as a candidate pocket embedded within a broader cavity-bearing region, not as a proven active site or experimentally validated ligand-binding interface. Likewise, although the current quantitative and structural evidence strongly supports an architecture-level and cavity-coupled loss-of-function interpretation, it does not yet establish the exact biochemical role of the disrupted region, the identity of any physiological substrate or ligand, or the specific molecular partners whose engagement may depend on this C-terminal module. These boundaries also apply to species transferability. The detailed mechanistic dissection was developed on the full-length rat Cwh43 scaffold, whereas supplementary inspection of the full-length human AlphaFold prediction together with InterPro-based annotation of the human C-terminal region supported the same high-level architectural conclusion that residue 533 lies within a domain-supported structured C-terminal region rather than a distal unstructured tail. The most justified level of transferability at present is therefore conservation of the internal-truncation logic at the module level, not identity of all residue-level features across species.

These boundaries and mechanistic alternatives help define the most informative next questions. By creating a premature termination codon (PTC), the residue-533 lesion may reduce transcript abundance through nonsense-mediated decay (NMD), while any residual truncated product, if produced, could additionally contribute through architecture-level disruption of the conserved C-terminal module described here. Accordingly, the present structure-guided analysis is most informative in defining the architectural consequences that would be expected from a truncated product, whereas future work will be needed to distinguish transcript-loss effects from residual truncated-protein effects rather than assuming a single-layer mechanism.

Important future directions therefore include whether the truncation perturbs specific partner-binding surfaces, alters protein-protein interaction organization, or affects possible dimeric or higher-order assembly states in membrane-associated contexts. More broadly, it remains important to determine whether the disrupted C-terminal structural unit contributes primarily to substrate engagement, complex assembly, membrane-proximal organization, or a combination of these functions. In this sense, the present study provides more than a structural annotation of a truncating variant: it establishes a practical mechanistic framework for future testing in rodent mutation models and related functional studies. Overall, the available evidence supports a model in which the disease-associated CWH43 residue-533 truncation disrupts a broader structurally integrated C-terminal cavity-bearing module rather than merely deleting a distal tail segment or a local pocket subset, thereby providing a more precise mechanistic bridge between human genetic association and predicted loss of function.

### Supplementary material note

Supplementary Fig. S1 provides extended support for the internal-truncation interpretation of residue 533, including the rat full-length framework, human full-length architectural confirmation, residue-533 truncation framing, and cross-species module-level transferability support. Supplementary Fig. S2 provides regional support across the C-terminal D3/Pfam-supported core, including conservation views and provisional hotspot-to-region linkage. Supplementary Fig. S3 provides expanded rat-side structural support for module-level disruption after residue-533 truncation, including five-model reproducibility, residue-class mapping, quantitative summaries, lightweight network support, deeper retained-side exemplars, and representative geometry measurements. Supplementary Table S1 summarizes residue-level structural integration features across rat D3, whereas Supplementary Table S2 summarizes the human variant interpretive framework used for Fig. 1d.

## Materials and methods

### Study design and overall analytical strategy

This study was designed to refine the mechanistic interpretation of the disease-associated human CWH43 variant c.1596del (ENST00000226432.9, MANE Select), annotated as p.Leu533Ter and referred to here mechanistically as the residue-533 truncation, by integrating structure prediction, domain-level annotation, conservation analysis, pocket/cavity mapping, cross-boundary contact analysis, and multi-model geometric reproducibility assessment. The central analytical question was whether residue 533 lies within a dispensable distal tail or within a structurally integrated C-terminal module whose internal organization is disrupted by truncation. All analyses were therefore organized around the full-length CWH43 model and the residue-533 truncation boundary.

### Protein sequence and structural models

The main structure-guided dissection in this study was performed using the full-length rat Cwh43 sequence and its associated AlphaFold-based structural framework, which was used as the reference model for mapping, visualization, pocket/cavity analysis, cross-boundary contact analysis, and representative residue-pair measurements. Five AlphaFold models from the rat prediction package were used to assess model-to-model reproducibility, with the highest-ranked model serving as the main reference structure for visualization and residue-level analysis. For display, the five AlphaFold models are denoted M0–M4, corresponding to model_0–model_4 in the rat prediction package. In parallel, the full-length human CWH43 AlphaFold prediction was inspected as an architecture-level confirmation resource to assess whether the disease-associated human CWH43 variant c.1596del (p.Leu533Ter) could be interpreted within the same high-level C-terminal structural context. This human-side confirmation was further supported by InterPro-based domain annotation of the human C-terminal region, which placed residue 533 within a domain-supported structured C-terminal region. Human-side confirmation was therefore used to support transferability of the internal-truncation interpretation but was not expanded into a second full pocket/cavity analysis pipeline.

### Operational domain framework

For mutation-centered interpretation, the protein was operationally partitioned into three modules: D1 (residues 1– 190), D2 (191–406), and D3 (407–699). This framework was adopted because the residue-533 truncation falls within the C-terminal D3 region and because comparative sequence and structural analyses converged on D3 as the most constrained C-terminal module in the current working architecture. The residue-533 truncation was therefore interpreted as an internal truncation within D3 rather than as removal of a distal tail segment. For the human full-length confirmation panels, the same residue-533-center interpretive logic was applied at the level of a domain-supported structured C-terminal architecture, without implying complete identity of all downstream residue-level features between species.

### Multiple-sequence alignment and conservation analysis

To evaluate evolutionary constraint across the C-terminal region, two multiple-sequence-alignment datasets were used: a broad taxonomic alignment spanning diverse homologous sequences and a mammalian-focused alignment emphasizing closer vertebrate conservation. Homologous C-terminal sequences were collected by BLAST and assembled into broad and mammalian-focused FASTA sets. Multiple-sequence alignment was performed using MAFFT (Katoh and Standley 2013) and Clustal Omega (Sievers et al. 2011) during iterative alignment optimization and cross-checking, and the resulting alignments were used for downstream comparative conservation analysis. Broad taxonomic and mammalian-focused alignments showed the same qualitative conservation pattern across the D3 region. These cross-species conservation patterns were also useful for evaluating transferability of the internal-truncation interpretation from the rat full-length framework to the human residue-533 variant context, although they were not used to claim strict residue-level identity across species.

### Pocket and cavity analysis

Surface-site analysis was performed at two complementary levels. First, the isolated D3 region (residues 407–699) was analyzed using a PrankWeb/P2Rank-based workflow (Krivak and Hoksza 2018; Jendele et al. 2019) to identify dominant candidate pocket features. Second, an independent cavity-mapping workflow based on DoGSiteScorer-style segmentation (Volkamer et al. 2012) was used to delineate broader cavity-associated structure and to classify core and extension components intersected by the truncation boundary.

### Cross-boundary contact and retained-side decoupling analysis

To evaluate whether the predicted structural effect of truncation extends beyond simple residue deletion, residue-level contacts spanning the residue-533 boundary were analyzed within D3. Contacts between residues retained N-terminal to the truncation site and residues deleted C-terminal to the truncation site were identified using heavy-atom distance criteria. Retained-side residues contacting deleted-side pocket-associated residues were then assessed for enrichment relative to the background retained D3 segment. This analysis was used to define a retained-but-contact-lost subset, namely residues that remain present after truncation but are predicted to lose deleted-side structural partners required for native local packing.

### Boundary-distance and local geometry analysis

To determine whether retained-side effects were confined to the immediate truncation edge or extended more deeply into D3, the positions of contact-linked retained residues were summarized relative to the residue-533 boundary. Representative retained/deleted residue pairs were then selected to illustrate both boundary-proximal and deeper cavity-associated coupling. For these pairs, minimum heavy-atom distances were measured across all five AlphaFold models and summarized as mean ± standard deviation and range. Side-chain centroid distances were also recorded as an auxiliary geometry measure for the representative-geometry summary. The main-text boundary-proximal representative pair was E526–N551, whereas W502–H548 was retained as a deeper retained-side exemplar for structural-consequence analysis. E466–H548 and P434–Y581 were retained as deeper supplementary exemplars.

### Lightweight network support

As a supplementary, non-primary analytical layer, a lightweight residue-contact graph was used to assess whether retained overlap-core residues were preferentially connected to deleted pocket-associated residues. This analysis was used only as supportive evidence and not as an independent pillar of the main structural claim. Network-derived results were therefore interpreted conservatively and mainly used to reinforce the broader view that retained-side decoupling is preferentially localized to cavity-associated structural features.

### Structural visualization and figure preparation

Structural visualization and figure preparation were performed using the reference AlphaFold model together with simplified residue-class overlays and module-level annotations. For Figure 1c, an online three-dimensional structure viewer was used to generate a full-length context inset and a D3-focused structural view, and final residue marking and panel assembly were adjusted during figure preparation to highlight the position of M533 within the folded D3 scaffold. Local structural inset views for the boundary-proximal exemplar (E526–N551; Fig. 2b) and the deeper retained-side exemplar (W502–H548; Fig. 3e) were generated from the reference AlphaFold model using the same three-dimensional visualization workflow, with residue-specific coloring and final panel assembly performed during figure preparation. Visualizations were designed to distinguish retained versus deleted segments, D3 versus non-D3 regions, and pocket/cavity-associated residues versus background structure. Multi-panel figures were organized so that the main text carried the most load-bearing mechanistic logic, whereas supplementary figures provided expanded geometry, reproducibility, and context views.

### Statistical analysis and interpretive framework

Simple descriptive summaries were used throughout for residue counts, retained/deleted proportions, and model-to-model geometric reproducibility. Where enrichment statistics were used, they served to assess whether retained-side contact-linked residues were disproportionately associated with cavity-related structural features relative to background composition. All statistical results were interpreted within a structure-guided inferential framework and were not used to claim biochemical validation. The strongest conclusions of the study were therefore kept at the module-level and architecture-level, namely that residue 533 lies within a structured C-terminal D3 module and that truncation is predicted to disrupt a broader cavity-bearing structural unit.

### Use of artificial intelligence-assisted tools

During manuscript preparation, the authors used ChatGPT and Claude for guidance on publicly available structural and sequence-analysis tools, manuscript editing, and consistency checks. All scientific content was reviewed and verified by the authors, who take full responsibility for the final manuscript.

## Figure legends

## Supplementary figure legends

**Supplementary Figure S1** Rat framework, human confirmation, and cross-species support for residue-533 internal truncation **a**, Rat full-length framework showing the D1–D3 organization, the position of residue 533, and the full-length scaffold used for the main structure-guided dissection. **b**, Rat C-terminal truncation boundary showing that the residue-533 boundary falls within the conserved C-terminal module rather than beyond it. **c**, Rat residue-533 truncation framing, using a D3-focused structural support view to indicate that residue 533 lies within the structured rat D3 segment. **d**, Human full-length structural confirmation showing that the human residue-533 position also falls within a domain-supported structured C-terminal region rather than a distal unstructured tail. **e**, Human residue-533 truncation framing in the full-length human context, illustrating retention of residues 1–532 and deletion of residues 533–699 within a domain-supported structured C-terminal region. **f**, Cross-species transferability support, summarizing rat and human architectural correspondence at the level of the structured C-terminal module. Transferability is supported at the module level rather than complete residue-wise identity.

**Supplementary Figure S2** Cross-species conservation and provisional hotspot prioritization across the C-terminal D3/Pfam-supported region **a**, Rat C-terminal D3/Pfam-supported region overview, showing the regional framework used for the support analyses in this figure and the position of the residue-533 boundary within the structured C-terminal support region. **b**, Cross-species conservation support across the D3/Pfam-defined core, using broad taxonomic and mammalian-focused alignment views to show that the C-terminal region analyzed in the rat scaffold is evolutionarily constrained across species. **c**, Rat-structure-based provisional hotspot prioritization, summarizing first-pass candidate surface classes within the structured C-terminal D3/Pfam-supported core. These classes serve as support-layer prioritization rather than definitive functional assignments. **d**, Hotspot-to-region linkage support, showing that the provisional hotspot classes summarized in panel c are linked to the same structured C-terminal D3/Pfam-supported region relative to the residue-533 boundary. This panel provides regional support only and is not intended as a variant-level interpretive layer.

Supplementary Figure S3 Expanded rat-side structural support for module-level disruption after residue-533 truncation a, Representative pair reproducibility across five rat AlphaFold models (M0–M4). b, Residue-class map across rat D3, providing the classification key used for panels c–e. c, Quantitative support for retained-side effects, summarizing overlap enrichment and boundary-distance spread beyond the immediate residue-533 boundary. d, Lightweight network support, showing a supportive network-derived signal rather than a primary mechanistic layer. e, Deeper retained-side exemplars, combining E466–H548 and P434–Y581 to illustrate additional retained/deleted residue pairs beyond the main-text exemplars and to support deeper cavity-associated structural coupling within D3. f, Representative geometry table summarizing five-model measurements for E526–N551, W502–H548, E466– H548, and P434–Y581. In panel f, minimum distance refers to the mean minimum heavy-atom distance, and side-chain centroid refers to the corresponding side-chain centroid distance across the five AlphaFold models.

## Statements and Declarations

### Author Contributions

Study conception and design were performed by Haowei Cao, Zhihan Yan, and Dejun Yang. Structural analysis, data integration, and figure organization were performed by Haowei Cao and Zhihan Yan. Jing Wang, Yue Liu, Zheng Wei and Yu Cheng contributed to data collection, analysis support, and manuscript discussion. Funding acquisition and supervision were provided by Haowei Cao, Shuqun Hu, and Dejun Yang. The first draft of the manuscript was written by Haowei Cao and Dejun Yang, and all authors commented on previous versions of the manuscript. All authors read and approved the final manuscript.

### Data Availability

The datasets and derived structural annotations generated and/or analyzed during the current study are available from the corresponding author on reasonable request. Source structural models and analysis-derived summary files used for this manuscript are described in the Supplementary Information.

### Ethics Approval

This manuscript reports a structure-guided computational analysis and did not involve new experiments in human participants or animals. Any previously published animal or human studies discussed for background or interpretation were conducted under the ethical approvals reported in the original publications.

### Consent to Participate

Not applicable.

### Consent for Publication

Not applicable.

