## Supplementary Figures for "Structure-guided reinterpretation of a disease-associated CWH43 residue-533 truncation identifies internal disruption of a conserved C-terminal module in idiopathic normal pressure hydrocephalus"

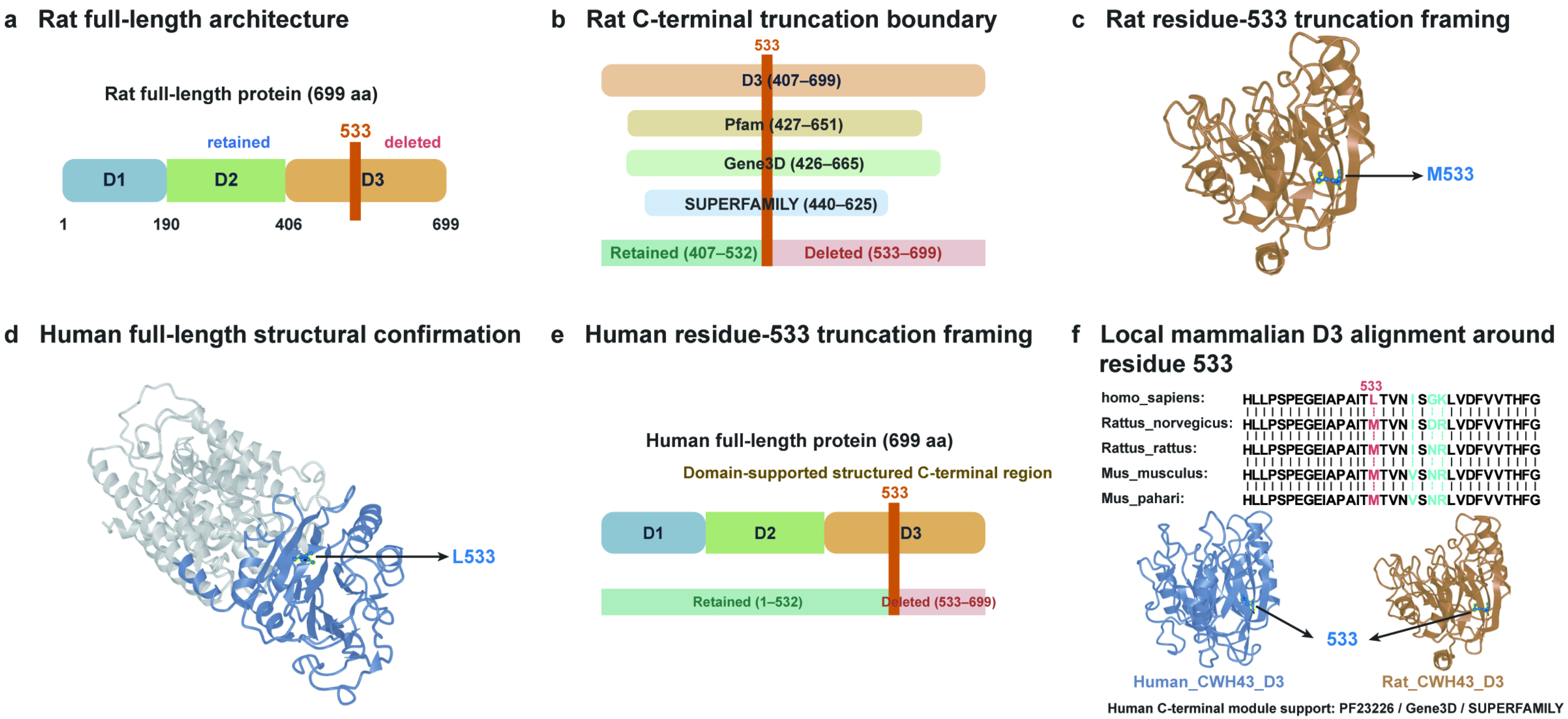
a Rat full-length architecture
Rat full-length protein (699 aa)
retained
533
deleted
D1
D2
D3
1
190
406
699
b Rat C-terminal truncation boundary
533
D3 (407–699)
Pfam (427–651)
Gene3D (426–665)
SUPERFAMILY (440–625)
Retained (407–532)
Deleted (533–699)
c Rat residue-533 truncation framing
M533
d Human full-length structural confirmation
L533
e Human residue-533 truncation framing
Human full-length protein (699 aa)
Domain-supported structured C-terminal region
533
D1
D2
D3
Retained (1–532)
Deleted (533–699)
f Local mammalian D3 alignment around residue 533
533
homo_sapiens: HLLPSPEGEIAPAITLTVN SGKLVDFVVTHFG
Rattus_norvegicus: HLLPSPEGEIAPAITMTVN SDRLVDFVVTHFG
Rattus_rattus: HLLPSPEGEIAPAITMTVN SNRLVDFVVTHFG
Mus_musculus: HLLPSPEGEIAPAITMTVNVSNRLVDFVVTHFG
Mus_pahari: HLLPSPEGEIAPAITMTVNVSNRLVDFVVTHFG
533
Human_CWH43_D3
Rat_CWH43_D3
Human C-terminal module support: PF23226 / Gene3D / SUPERFAMILY

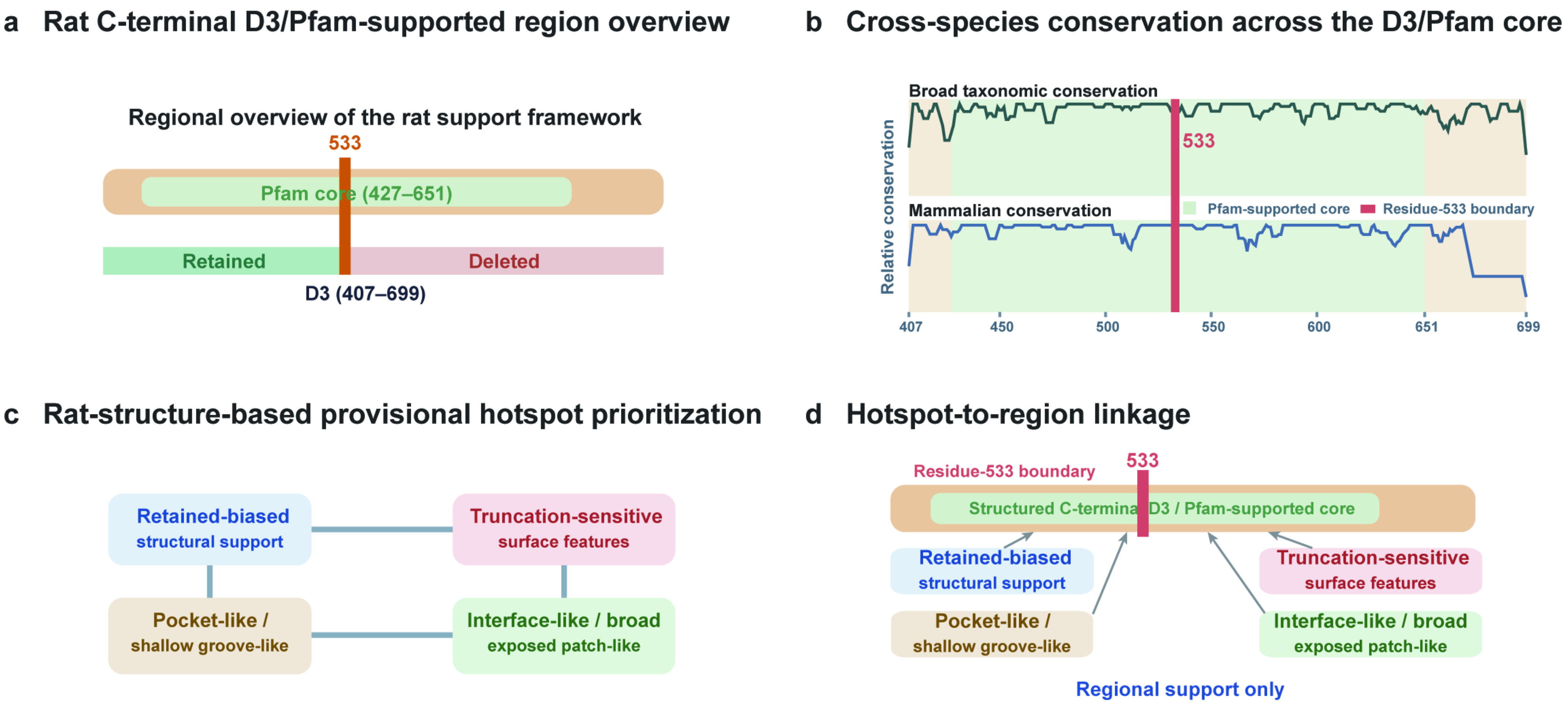
a Rat C-terminal D3/Pfam-supported region overview
Regional overview of the rat support framework
533
Pfam core (427–651)
Retained
Deleted
D3 (407–699)
b Cross-species conservation across the D3/Pfam core
Broad taxonomic conservation
533
Mammalian conservation
Pfam-supported core
Residue-533 boundary
Relative conservation
407
450
500
550
600
651
699
c Rat-structure-based provisional hotspot prioritization
Retained-biased
structural support
Truncation-sensitive
surface features
Pocket-like /
shallow groove-like
Interface-like / broad
exposed patch-like
d Hotspot-to-region linkage
Residue-533 boundary
533
Structured C-terminal D3 / Pfam-supported core
Retained-biased
structural support
Truncation-sensitive
surface features
Pocket-like /
shallow groove-like
Interface-like / broad
exposed patch-like
Regional support only

**a** Representative pair reproducibility

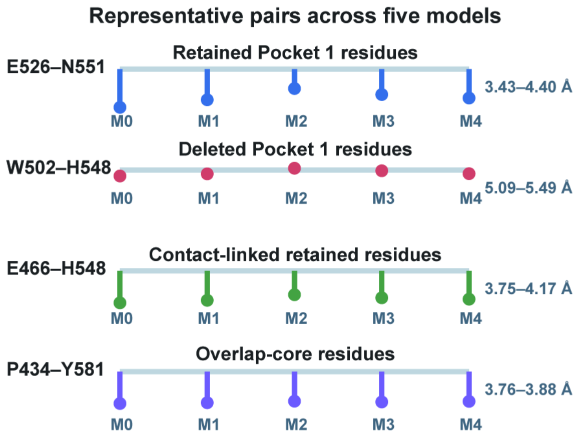


**b** Residue-class mapping across D3

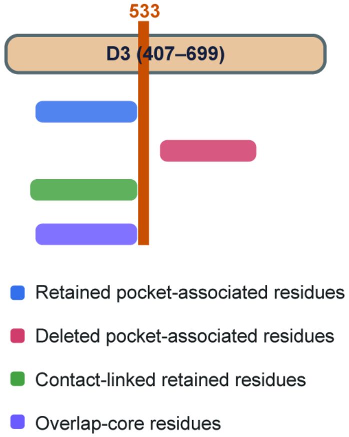


**c** Quantitative support for retained-side effects

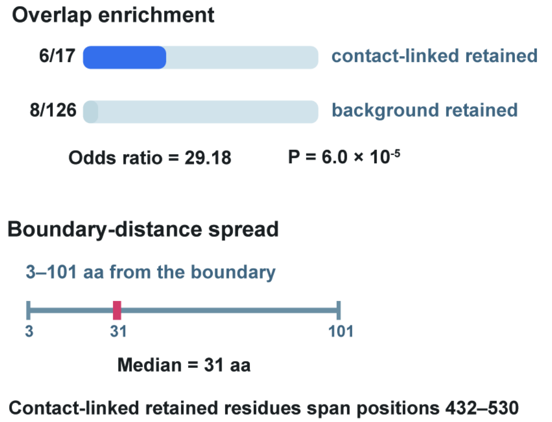


**d** Lightweight network support

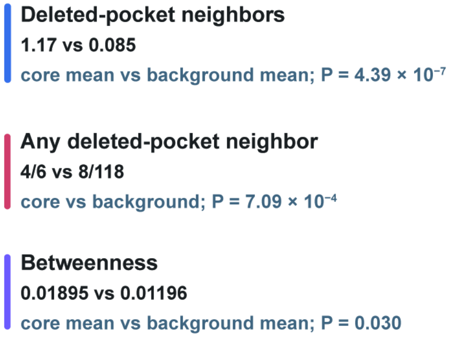


**e** Deeper supplementary exemplars

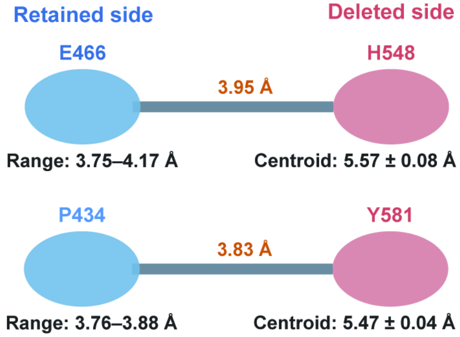


**f** Representative geometry table

| Pair | Mean ± SD (Å) | Range (Å) | Side-chain centroid (Å) | Recommended role |
|---|---|---|---|---|
| E526–N551 | 3.93 ± 0.35 | 3.43–4.40 | 5.66 ± 0.13 | Boundary-proximal exemplar |
| W502–H548 | 5.27 ± 0.16 | 5.09–5.49 | 8.11 ± 0.12 | Deeper retained-side exemplar |
| E466–H548 | 3.95 ± 0.16 | 3.75–4.17 | 5.57 ± 0.08 | Deeper supplementary exemplar |
| P434–Y581 | 3.83 ± 0.05 | 3.76–3.88 | 5.47 ± 0.04 | Very deep supplementary exemplar |
